# Enterococcal quorum-controlled protease alters phage infection

**DOI:** 10.1101/2024.05.10.593607

**Authors:** Emma K. Sheriff, Fernanda Salvato, Shelby E. Andersen, Anushila Chatterjee, Manuel Kleiner, Breck A. Duerkop

**Author notes:** Correspondence: Breck A. Duerkop, 1.

## Abstract

Increased prevalence of multidrug resistant bacterial infections has sparked interest in alternative antimicrobials, including bacteriophages (phages). Limited understanding of the phage infection process hampers our ability to utilize phages to their full therapeutic potential. To understand phage infection dynamics we performed proteomics on *Enterococcus faecalis* infected with the phage VPE25. We discovered numerous uncharacterized phage proteins are produced during phage infection of *Enterococcus faecalis*. Additionally, we identified hundreds of changes in bacterial protein abundances during infection. One such protein, enterococcal gelatinase (GelE), an *fsr* quorum sensing regulated protease involved in biofilm formation and virulence, was reduced during VPE25 infection. Plaque assays showed that mutation of either the *fsrA* or *gelE* resulted in plaques with a “halo” morphology and significantly larger diameters, suggesting decreased protection from phage infection. GelE-associated protection during phage infection is dependent on the murein hydrolase regulator LrgA and antiholin-like protein LrgB, whose expression have been shown to be regulated by GelE. Our work may be leveraged in the development of phage therapies that can modulate the production of GelE thereby altering biofilm formation and decreasing *E. faecalis* virulence.

## INTRODUCTION

*Enterococcus faecalis* is a Gram-positive bacterium and a member of the gut microbiota of diverse animals, including humans (1–3). Following prolonged antibiotic therapy, *E. faecalis* can outgrow other members of the microbiota and disseminate to the bloodstream, leading to life threating disease such as sepsis and endocarditis (4–7). Additionally, *E. faecalis* is a common cause of healthcare-associated infections (8–10). Treatment of enterococcal infections is complicated by the increasing prevalence of multi-drug resistant (MDR) *E. faecalis* strains, including those resistant to “last-resort” antibiotics (11–14). With the ongoing antibiotic discovery gap and rising incidence of MDR infections, it is estimated that over 10 million individuals may die of antibiotic resistant infections per year by 2050, nearly ten times the current yearly mortality (15, 16). Therefore, the development of innovative antimicrobial therapies is crucial to combating antibiotic resistant bacteria.

Bacteriophages (phages) are viruses that infect and kill bacteria. They have reemerged as potential therapeutics due to their diversity and abundance in nature, making them readily available for medical applications (17, 18). Despite the discovery of phages over 100 years ago, we know little about the function of most phage-encoded genes (19, 20). Additionally, we have only a rudimentary understanding of how phages interact with their target bacteria, particularly in non-model hosts (21). Understanding these fundamental aspects of phage biology is an important milestone toward the development of phages as antimicrobials.

During infection, phages co-opt host cellular processes to support their genome replication and translation of proteins responsible for virion assembly (22, 23). One way that phages are known to influence the bacterial response to infection is by modulating the cell density-dependent transcriptional program known as quorum sensing (24–27). Phage-mediated changes in quorum sensing gene regulation within a bacterial population can create an environment that is more permissible for infection (24, 25). For example, changes in quorum sensing can result in the modification of wall teichoic acids, a major phage receptor, which contributes to phage infectivity (28, 29). In *E. faecalis*, high levels of the quorum-sensing peptide AI-2 are associated with release of prophages, possibly resulting in the transfer of virulence genes (26). Additionally, *E. faecalis* quorum sensing itself is interrupted during phage infection, with decreased transcription of quorum-sensing-induced genes and increased transcription of quorum-sensing-repressed genes (30).

Quorum sensing in *E. faecalis* is controlled by the Fsr system (27, 31). The transcription factor FsrA positively regulates three operons in response to bacterial density, as sensed by the extracellular abundance of the gelatinase biosynthesis-activating pheromone (GBAP) (32). The first of these is the *fsrBDC* quorum-sensing locus, which encodes the machinery for the quorum-sensing system. This includes FsrC, the transmembrane sensor-transmitter; FsrD, a precursor of GBAP; and FsrB, which processes the FsrD propeptide into GBAP (32–34). The second operon controlled by FsrA encodes the enterocin EntV, which has activity against Gram-positive bacteria, fungi, and *Drosophila* (35–37). The final operon regulated by Fsr quorum-sensing contains the genes *gelE* and *sprE* (31, 35). GelE, or gelatinase, is a secreted metalloprotease frequently associated with virulence in enterococci and is vital for initiation of biofilm formation (38–42). SprE is a serine protease also associated with virulence (39, 40, 43). In addition to its roles in virulence, GelE plays a role in the post-processing of SprE and EntV (44, 45). Additionally, GelE and SprE have both been shown to be involved in the post-processing of the autolysin AtlA, although the role of SprE in this context has been contested (41, 46).

In this paper, we used a proteomic approach to explore the interaction between the enterococcal phage VPE25 and its *E. faecalis* host. We found time-dependent trends in phage protein production and identified correlations between gene expression and protein abundance levels during infection. Investigation of the changes in bacterial protein abundance during phage predation revealed large scale trends in the abundance of a wide variety of proteins. We show that decreased levels of the quorum-sensing-regulated protein GelE, and subsequent downstream changes in the production of the murein hydrolase modulator LrgA and antiholin-like protein LrgB, influence the outcome of phage infection.

## MATERIALS AND METHODS

### Bacterial strains, bacteriophages, and plasmids

All bacterial strains used in this study have been previously published and are detailed in Supplemental Table 1 (36, 47–53). Transposon mutants selected from an arrayed library were confirmed using PCR. Phages, plasmids, and primers used in this study are also detailed in Supplemental Table 1 (54–58). *E. faecalis* strains were grown in Todd-Hewitt broth (THB) with aeration or on THB agar plates at 37°C. Complementation and empty vectors were selected in *E. faecalis* strains using 15 μg/mL chloramphenicol as appropriate.

### RNA-seq analysis

A previously published RNA-seq dataset, generated in a parallel experiment, was reanalyzed for phage transcript abundances (EMBL-EBI ArrayExpress database, accession number E-MTAB-8546) (30, 59). RNA sequencing reads were analyzed for quality using FastQC v0.12.0 (60). Reads were then mapped to the VPE25 open reading frames (ORFs) by Salmon v1.10.2 (61). The output read quantities, as reported in the nascent quant.sf file, were subject to relative abundance calculations by dividing the transcripts per million (TPM) for a given ORF by the total number of TPMs for a sample. ORFs were ranked by their abundance for each timepoint, and this ranking was used to inform heatmap rings in Figure 1A, generated using the R package Circlize (62).

**Figure 1:**
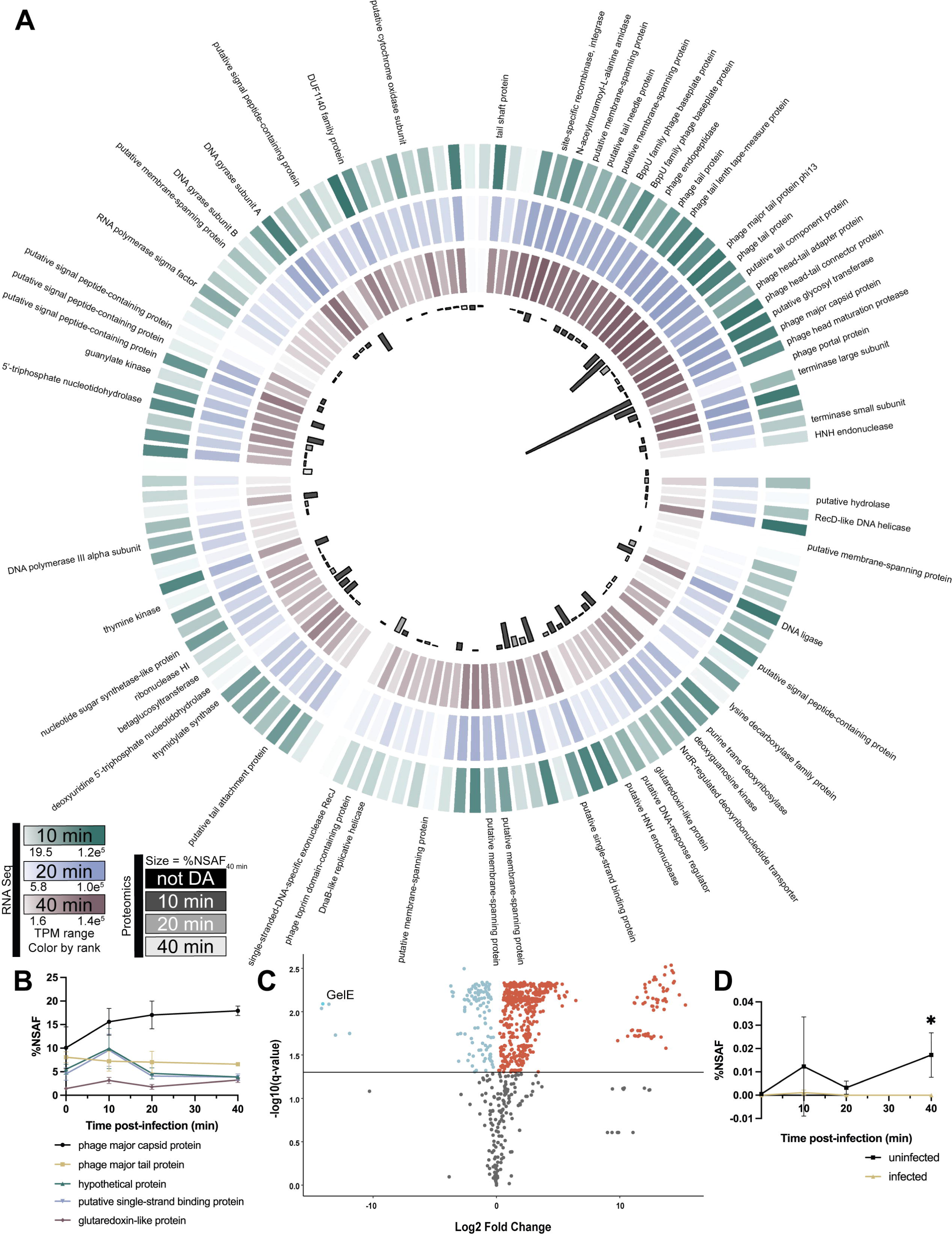
Phage infection of *E. faecalis* leads to diverse changes in protein expression. **(A)** Combined transcriptomic and proteomic data for phage VPE25. Each column represents one phage gene, with available gene annotations listed alongside corresponding columns. The outer three rings represent average transcripts per million (TPM) colored by rank at 10, 20, and 40 minutes post-infection. The innermost ring represents the percentage normalized spectral abundance factors (%NSAF) of each corresponding protein at 40 minutes post-infection. %NSAF abundance bars are color coded by the timepoint at which they are first identified as significantly differentially abundant (DA). **(B)** %NSAF values of the five most abundant phage proteins at 40 minutes post-infection are charted over the course of infection. Points represent an average of at least three biological replicates. Error bars represent standard deviation. **(C)** Volcano plot of bacterial proteins differentially abundant at 40 minutes post-infection between the infected and uninfected samples. Log2-transformed fold-change between the two samples is shown on the x-axis. Negative log10-transformed q-value is shown on the y-axis with a line at a significance cut-off of q < 0.05. Points colored in blue are significantly underrepresented in the infected sample when compared to the uninfected sample. Points in red are significantly overrepresented. Proteins with a q-value of zero were excluded from the chart. **(D)** %NSAF of GelE over time in both the infected and uninfected samples. Points represent an average of four biological replicates. Significance was determined using student’s t-tests corrected for multiple hypothesis testing using a permutation-based false discovery rate of 5% (see Materials and Methods: Protein quantifications and statistical analyses). Error bars represent standard deviation. * FDR ≤ 5%.

### Preparation of samples for proteomics

Subcultures of *E. faecalis* overnight cultures were infected with VPE25 as described previously (30). Four samples of 4 mL each were taken at 0, 10, 20, and 40 minutes after VPE25 treatment and pelleted. Pelleted samples were resuspended in 300 µL of SDT-lysis buffer (4% (w/v) SDS, 100 mM Tris-HCl, 0.1LM DTT). Cells were lysed by bead-beating using a Bead Ruptor Elite (OMNI) with Matrix Z beads (MP Biomedicals) for two cycles of 45 s at 6 m/s. Samples were incubated at 95°C for 10 min. Tryptic digests of protein extracts were prepared following the filter-aided sample preparation (FASP) protocol described previously, with minor modifications as described in Kleiner et al. (63, 64). Lysate was not cleared by centrifugation after boiling the sample in lysis buffer. The whole lysate was loaded onto the filter units used for the FASP procedure. Centrifugation times were reduced to 20 minutes as compared to Kleiner et al. (64). Peptide concentrations were determined with the Pierce Micro BCA assay (Thermo Scientific) using an Epoch2 microplate reader (Biotek) following the manufacturer’s instructions.

### LC-MS/MS

All samples were analyzed by 1D-LC-MS/MS as described previously, with the modification that a 75 cm analytical column and a 140-minute long gradient were used (65). For each sample run, 400 ng peptide were loaded with an UltiMateTM 3000 RSLCnano Liquid Chromatograph (Thermo Fisher Scientific) in loading solvent A (2% acetonitrile, 0.05% trifluoroacetic acid) onto a 5 mm, 300 µm ID C18 Acclaim® PepMap100 pre-column (Thermo Fisher Scientific). Separation of peptides on the analytical column (75 cm x 75 µm analytical EASY-Spray column packed with PepMap RSLC C18, 2 µm material, Thermo Fisher Scientific) was achieved at a flow rate of 300 nl min!Zl1 using a 140 min gradient going from 95% buffer A (0.1% formic acid) to 31 % buffer B (0.1% formic acid, 80% acetonitrile) in 102 min, then to 50% B in 18 min, to 99% B in 1 min and ending with 99% B. The analytical column was heated to 60°C and was connected to a Q Exactive HF hybrid quadrupole-Orbitrap mass spectrometer (Thermo Fisher Scientific) via an Easy-Spray source. Eluting peptides were ionized via electrospray ionization (ESI). Carryover was reduced by one wash run (injection of 20 µl acetonitrile, 99% eluent B) between samples. Full scans were acquired in the Orbitrap at 60,000 resolution. The 15 most abundant precursor ions were selected for fragmentation and MS/MS scans were acquired at 15,000 resolution. The mass (m/z) 445.12003 was used as lock mass. Ions with charge state +1 were excluded from MS/MS analysis. Dynamic exclusion was set to 18 s. Roughly 120,000 MS/MS spectra were acquired per sample.

### Protein identification

A database containing protein sequences from *E. faecalis* (RefSeq accession: NC_017316.1) and the phage VPE25 (GenBank accession: LT615366.1), both downloaded from NCBI, were used (66, 67). Sequences of common laboratory contaminants were included by appending the cRAP protein sequence database (http://www.thegpm.org/crap/). The final database contained 2,893 protein sequences. Searches of the MS/MS spectra against this database were performed with the Sequest HT node in Proteome Discoverer version 2.2.0.388 (Thermo Fisher Scientific) as previously described (68). Only proteins identified with medium or high confidence were retained resulting in an overall false discovery rate of <5%.

### Protein quantification and statistical analyses

For quantification of bacterial proteins, normalized spectral abundance factors (NSAFs) were calculated and multiplied by 100% to obtain relative protein abundance (69). The NSAF values were loaded into Perseus version 1.6.2.3 and log_2_ transformed (70). Only proteins with valid values in four replicates of at least one treatment were considered for further analysis. Missing values were replaced by a constant number that was lower than any value across the experiment. Differentially abundant proteins between conditions were calculated using student’s t-tests corrected for multiple hypothesis testing using a permutation-based false discovery rate of 5%.

As the number of phage proteins is small and their relative abundances are heavily influenced by the overall abundance of the bacterial proteins in the cultures, we used a centered-log ratio (CLR) transformation for more robust analysis of compositional data (71). The CLR was calculated for all proteins, both phage and bacterial. CLR values for phage proteins were loaded into Perseus 1.6.2.3 where t-tests were performed as described above (70). Since the spectral counts at t0 (0 minutes post-infection) were much lower than the other time points, only proteins with 70% of total values across conditions with more than 20 PSMs were considered for comparisons with t0. For comparisons between the 10, 20, and 40 minutes post-infection time points, no PSM cut-off was employed. Replacement of missing values and t-tests were performed as described above.

### Plaque assays

One mL of overnight bacterial culture was pelleted at 21,000 ×Lg for 1 minute and resuspended in 2 mL SM-plus phage buffer (100 mM NaCl, 50 mM Tris-HCl, 8 mM MgSO_4_, 5 mM CaCl_2_ [pH 7.4]) (55). Approximately fifteen plaque forming units (PFU) of phage were added to 120 μL of resuspended bacterial culture and incubated statically at room temperature for 5 minutes. Following incubation, 5 mL of molten THB top agar (0.35% agar, unless otherwise noted) supplemented with 10mM MgSO_4_ was added to the suspension and poured over a 1.5% THB agar plate supplemented with 10 mM MgSO_4_. Plates were incubated upright at 37°C for 24 hours unless otherwise noted. Photos of plates were taken on an iPhone 12, and plaque diameters were measured using ImageJ v1.53t (72). A standard ruler was included in each image to determine scale.

## RESULTS

### Assessment of VPE25 transcript and protein abundances during *E. faecalis* infection

To understand global protein regulation and abundances during phage infection of *E. faecalis*, we infected *E. faecalis* OG1RF with the siphophage VPE25 (55). Mid-log cultures of *E. faecalis* were infected with VPE25 at a multiplicity of infection (MOI) of 10. Samples were taken at 0, 10, 20, and 40 minutes post-infection and subjected to LC-MS/MS-based proteomics analyses. We compared the phage-encoded protein abundances to a previously generated RNA-Seq dataset during VPE25 infection of *E. faecalis* OG1RF, where the experimental parameters were identical (30). From the RNAseq, we identified 131 phage transcripts throughout infection (Supplemental Table 2). Proteomic analysis identified 93 of those transcripts as proteins (Supplemental Table 3).

Figure 1A shows the VPE25 transcript and protein abundances throughout infection. Few proteins were detected at the initiation of infection, and it is likely that those detected were either present in the virion or rapidly expressed following DNA entry. Many replication proteins, such as DNA polymerase and DNA helicases, were significantly differentially abundant (DA) after only 10 minutes post-infection (Supplemental Table 3). By 20 minutes post-infection, we saw increases in abundance of additional replication machinery, such as DNA gyrase subunits A and B. Virion components, primarily proteins involved in assembly of the tail, were also DA at this time, as was the phage lysin. Final virion components, including the portal protein and the head maturation protease, were DA by 40 minutes post-infection. Notably, the most abundant phage protein at each timepoint was the major capsid protein, whose relative abundance continued to increase throughout the course of infection (Figure 1B, Supplemental Table 3). The major capsid transcript levels reached the top ten most abundant phage transcripts at the 10-minute timepoint and remained among the most abundant transcripts throughout infection (Supplemental Table 2).

A high number of detected phage transcripts and proteins were annotated as genes of unknown function. Of the 133 putative genes in the VPE25 genome, more than 60% are hypothetical. Many of these uncharacterized proteins were expressed at a level equal to or above proteins with characterized functions. For example, SCO93409.1 is an uncharacterized phage protein whose abundance at 40 minutes was third only to the major capsid and major tail proteins (Figure 1B). *In silico* analysis of these hypothetical proteins failed to provide insight on their potential function, with more than half having no conserved domains identified via InterProScan (73). Together these data show that the majority of VPE25 proteins are expressed during infection, many of which are proteins of unknown function.

### *E. faecalis* responds to VPE25 infection by altering its gene expression and protein abundances

Our understanding of bacterial gene expression levels in response to phage infection and how this corresponds to protein abundances is limited. While transcript data is often used as a stand-in for shifts in proteomic abundances, comparative analyses show that these are often not fully concordant (74, 75). In fact, many of the most differentially abundant bacterial proteins in our analysis are relatively unchanged at the transcript level when compared to our previously published RNA-Seq dataset (Table 1, Supplemental Table 4) (30).

**Table 1:** High and low differential abundance ratio (DAR) proteins at 40 minutes^1^.

| Gene Name | Description | DAR<sub>40</sub> | Recovered from transposon library? |
| --- | --- | --- | --- |
| **High DAR<sub>40</sub> Genes** |  |  |  |
| *secY* | Preprotein translocase subunit SecY | 86.97 | No, not present |
| *OG1RF_11743* | TetR/AcrR family transcriptional regulator | 52.26 | Yes |
| *OG1RF_11417* | Pyridoxal phosphate-dependent aminotransferase | 47.66 | Yes |
| *OG1RF_10804* | ATP-binding protein | 45.55 | Yes |
| *OG1RF_12485* | Alkaline phosphatase family protein | 40.55 | Yes |
| *phoU* | Phosphate signaling complex protein | 40.16 | Yes |
| *OG1RF_10823* | AAA family ATPase | 34.49 | Yes |
| *OG1RF_11853* | Homoserine dehydrogenase | 33.26 | Yes |
| *yidC* | Membrane protein insertase | 29.87 | Yes |
| *bgsB* | Glycosyltransferase family 4 protein | 27.65 | Yes |
| *rsmG* | 16S rRNA (guanine(527)-N(7))-methyltransferase | 27.43 | Yes |
| *OG1RF_11964* | Mur ligase family protein | 27.43 | No, not present |
| *citD* | Citrate lyase acyl carrier protein | 24.39 | No, not present |
| *OG1RF_11639* | DUF3013 family protein | 23.06 | Yes |
| *dltA* | D-alanine—poly(phosphoribitol) ligase subunit | 22.09 | Yes |
| *OG1RF_11645* | Class I SAM-dependent methyltransferase | 21.34 | No, not present |
| *OG1RF_10575* | Alanine ligase | 21.07 | No, not present |
| *OG1RF_12200* | NAD(P)/FAD-dependent oxidoreductase | 21.05 | No, not present |
| *OG1RF_12473* | tRNA-dihydrouridine synthase | 20.22 | Yes |
| *OG1RF_10734* | RNA-binding protein | 20.03 | Yes |
| **Low DAR<sub>40</sub> Genes** |  |  |  |
| *gelE* | M4 family metallopeptidase coccolysin | 0.00 | Yes |
| *OG1RF_10381* | Lipoate—protein ligase | 0.00 | Yes |
| *OG1RF_10282* | S-methyltransferase | 0.00 | Yes |
| *msrA* | Peptide-methionine (S)-S-oxide reductase | 0.00 | No, not present |
| *OG1RF_11885* | DUF2129 domain-containing protein | 0.00 | Yes |
| *OG1RF_10952* | CsbD family protein | 0.08 | No, not present |
| *OG1RF_11211* | V-type ATPase subunit | 0.08 | Yes |
| *OG1RF_10532* | DUF1700 domain-containing protein | 0.09 | Yes |
| *OG1RF_11274* | Nucleotide pyrophosphohydrolase | 0.09 | No, not present |
| *OG1RF_10750* | PTS sugar transporter subunit IIB | 0.11 | No, not present |
| *opp2D* | ABC transporter ATP-binding protein | 0.12 | Yes |
| *OG1RF_10751* | PTS lactose/cellobiose transporter subunit IIA | 0.13 | No, not present |
| *OG1RF_11463* | PspC domain-containing protein | 0.16 | No, not present |
| *OG1RF_10660* | AAA family ATPase | 0.16 | Yes |
| *OG1RF_11909* | Hypothetical protein | 0.17 | Yes |
| *OG1RF_12106* | MBL fold metallo-hydrolase | 0.17 | Yes |
| *atpF* | F0F1 ATP synthase subunit B | 0.17 | No, not present |
| *OG1RF_11402* | LapA family protein | 0.17 | No, not present |
| *OG1RF_11033* | LTA synthase family protein | 0.17 | No, not present |
| *OG1RF_10795* | Hypothetical protein | 0.19 | Yes |

<sup>1</sup> The differential abundance ratio at 40 minutes post-infection was calculated using the following formula: DAR<sub>40</sub> = (average %NSAF in phage infected sample at 40 minutes) / (average %NSAF in uninfected sample at 40 minutes)

Of the 2,647 genes in the *E. faecalis* OG1RF genome, 2,550 were detected as transcripts (30). Of those transcripts, 1,225 proteins were detected using proteomics. A total of 680 proteins were differentially abundant (DA) between infected and uninfected *E. faecalis* by 40 minutes (Figure 1C, Supplemental Table 4). There was no statistical difference in bacterial protein relative abundances at the 10-minute timepoint, and at 20 minutes post-infection only 6 proteins were differentially abundant (Supplemental Figure 1A). The limited differences in protein abundances at the earlier timepoints reflect a lag time between the transcriptional response, which begins by the 10-minute timepoint, and the time it takes to translate those transcripts into proteins (30). All of the DA proteins at 20 minutes remained DA at the 40-minute timepoint with the exception of OG1RF_12042, a CYTH domain-containing protein.

The differential abundance ratio, or DAR, was calculated as the relative change in abundance of a protein between the infected and uninfected samples at each timepoint. The DAR at 40 minutes (DAR_40_) was used to identify potential key effectors of bacteria-phage interactions during infection (Table 1). Proteins with high DAR values, such as SecY, are at an increased level during phage infection, while proteins with low DAR values, like AtpF, are found in decreased abundance during phage infection. This up- and down-regulation of protein abundances can be attributed to the bacterium responding to phage infection or when phage-encoded proteins hijack bacterial host processes leading to changes in protein abundances. To determine which, if any, of these proteins may be integral to phage virulence during infection, we used mutants from an arrayed *E. faecalis* OG1RF transposon library and screened them for changes in phage infectivity (48). Mutants selected represent available transposon insertions in genes whose proteins were most over- or underrepresented at 40 minutes as indicated by DAR (Table 1). While no differences were seen in the number of viable PFU on each strain tested (Supplemental Figure 1B), a change in plaque morphology was noted on the *gelE*-Tn mutant, a protein whose expression is undetected at 40 minutes (Figure 1D).

### *E. faecalis* gelatinase alters phage infection

VPE25 forms clear plaques with a well-defined border approximately 1 mm in diameter on *E. faecalis* OG1RF. However, after 24 hours of plaque formation on an in-frame *gelE* deletion strain (OG1RFΔ*gelE*), VPE25 forms a central zone of clearing surrounded by a large turbid ring which more than doubles the overall diameter (Figure 2A). This phenotype suggests that when GelE is absent, bacterial cells are more susceptible to phage infection associated factors. This “halo” morphology is rescued through the addition of *gelE* on a constitutive expression vector (pLZ12A::*gelE*) (Figures 2B, 2C). The *gelE* gene encodes the enterococcal gelatinase, a secreted protease that targets a variety of substrates and is regulated by the Fsr quorum-sensing system (31, 35, 44, 45, 76–80). GelE is one of five proteins, which became undetectable in the phage-infected sample by 40 minutes post-infection and thus has a DAR of 0, supporting our previous finding that the transcription of quorum-sensing regulated genes, including *gelE*, are significantly lower during phage infection (Table 1, Supplemental Table 4) (30).

**Figure 2:**
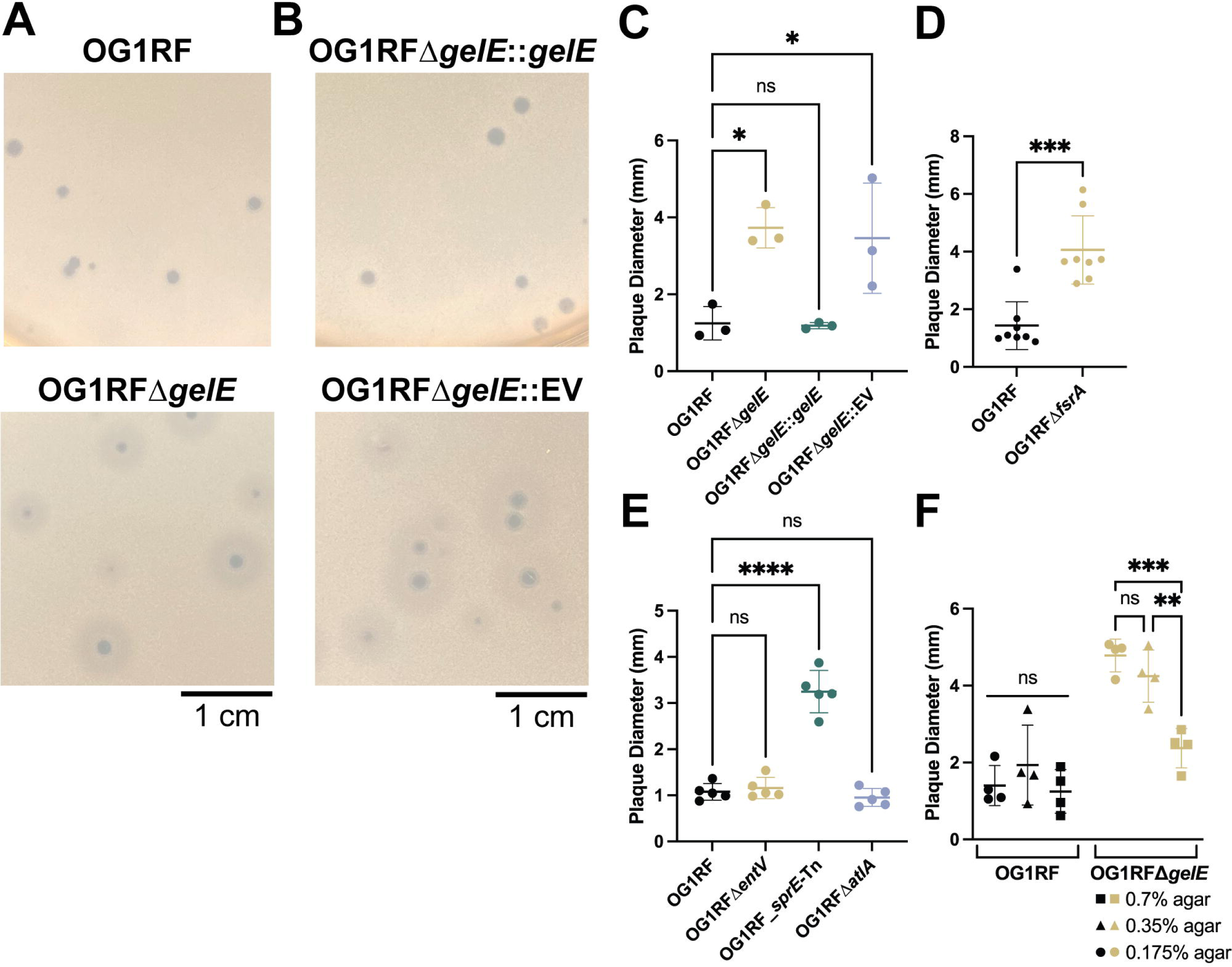
Phage VPE25 plaque morphology is altered in the absence of *gelE*. **(A)** VPE25 plaques on an OG1RF top agar overlay are small with distinct edges, while plaques on an OG1RFΔ*gelE* overlay are larger and have a halo phenotype. **(B)** Complementation of OG1RFΔ*gelE* with *gelE* on a plasmid (OG1RFΔ*gelE*::*gelE*) restores the OG1RF phenotype, while addition of the empty vector (OG1RFΔ*gelE*::EV) has no effect. **(C)** Average plaque diameters are significantly larger in OG1RFΔ*gelE* and OG1RFΔ*gelE*::EV than in OG1RF or OG1RFΔ*gelE*::*gelE*. Data represents average plaque diameter from three biological replicates. **(D)** VPE25 plaques on OG1RFΔ*fsrA* are significantly larger than on OG1RF. Data represents average plaque diameter from eight biological replicates. Significance determined using unpaired t test. **(E)** Average plaque diameters are significantly larger than wild type in overlays of *sprE*-Tn when compared to OG1RF. Data represents average plaque diameter from five biological replicates. **(F)** Plaque diameter on OG1RFΔ*gelE* varies with concentration of agar in top agar overlay. Data represents average plaque diameter from four biological replicates. Significance determined using two-way ANOVA with multiple comparisons. **(C, E)** Significance determined using one-way ANOVA with multiple comparisons. **(C, D, E, F)** Error bars represent standard deviation. **** p ≤ 0.0001, *** 0.0001 < p ≤ 0.001, ** 0.001 < p ≤ 0.01, * 0.01 < p ≤ 0.05, ns = not significant.

To further investigate this phenotype, we assessed phage plaque morphology on *E. faecalis* OG1RFΔ*fsrA*, a strain lacking the quorum-sensing master regulator *fsrA*. Again, plaques showed a halo morphology, and plaque diameter was significantly larger than wildtype (Figure 2D). GelE cleaves the target proteins EntV and SprE, which are both under the control of FsrA, as well as AtlA, an autolysin that is involved in biofilm formation (31, 36, 37, 41, 44–46). We tested deletion mutants of *entV*, *sprE*, and *atlA* for plaque morphology during VPE25 infection. Deletion of *entV* or *atlA* showed no change in plaque morphology or size, but a *sprE* transposon mutant strain formed haloed plaques similar to OG1RFΔ*gelE* and OG1RFΔ*fsrA* (Figure 2E). GelE-dependent changes in the viability of extracellular virions and ability of VPE25 to adsorb were also tested; however, we observed no significant differences regardless of the presence or absence of GelE (Supplemental Figures 2B, 2C).

The top agar overlay plaque assay supports the diffusion of both phage and secreted factors, such as GelE, through the top agar. We varied the agar density to look for changes in plaque size to determine if this phenotype is associated with diffusion. Plaque assays were performed using either double (0.7% agar) or half (0.175% agar) the normal concentration of soft agar in the overlay, while all other parameters remained constant. As the agar concentration within the overlay increased, the size of VPE25 plaques and their halos on OG1RFΔ*gelE* decreased, suggesting that a diffusible factor is indeed responsible for this phenotype (Figure 2F). When wild type OG1RF is infected with VPE25, the average plaque diameter does not change regardless of the agar concentration.

### Diverse *E. faecalis* bacterial strains and phages show altered infectivity that is dependent on GelE

To determine if the haloed plaque morphology was unique to *E. faecalis* OG1RF and VPE25, we tested VPE25 plaque morphology on *E. faecalis* V583, a genetically distinct vancomycin-resistant isolate (47). In a *gelE* mutant of V583 (V583Δ*gelE*), VPE25 plaques develop a similar halo morphology, and the plaque diameter is significantly larger when compared to wild type cells (Figure 3A). However, unlike OG1RFΔ*gelE*, the halo morphology is not well-observed at 24 hours and is only prominent in the V583 background after 48 hours (Supplemental Figure 3A). The halos observed in V583Δ*gelE* are also present in a double deletion of both *gelE* and *sprE* (V583Δ*gelE*Δ*sprE*), but not present in the *sprE* deletion alone (V583Δ*sprE*) (Figure 3A).

**Figure 3:**
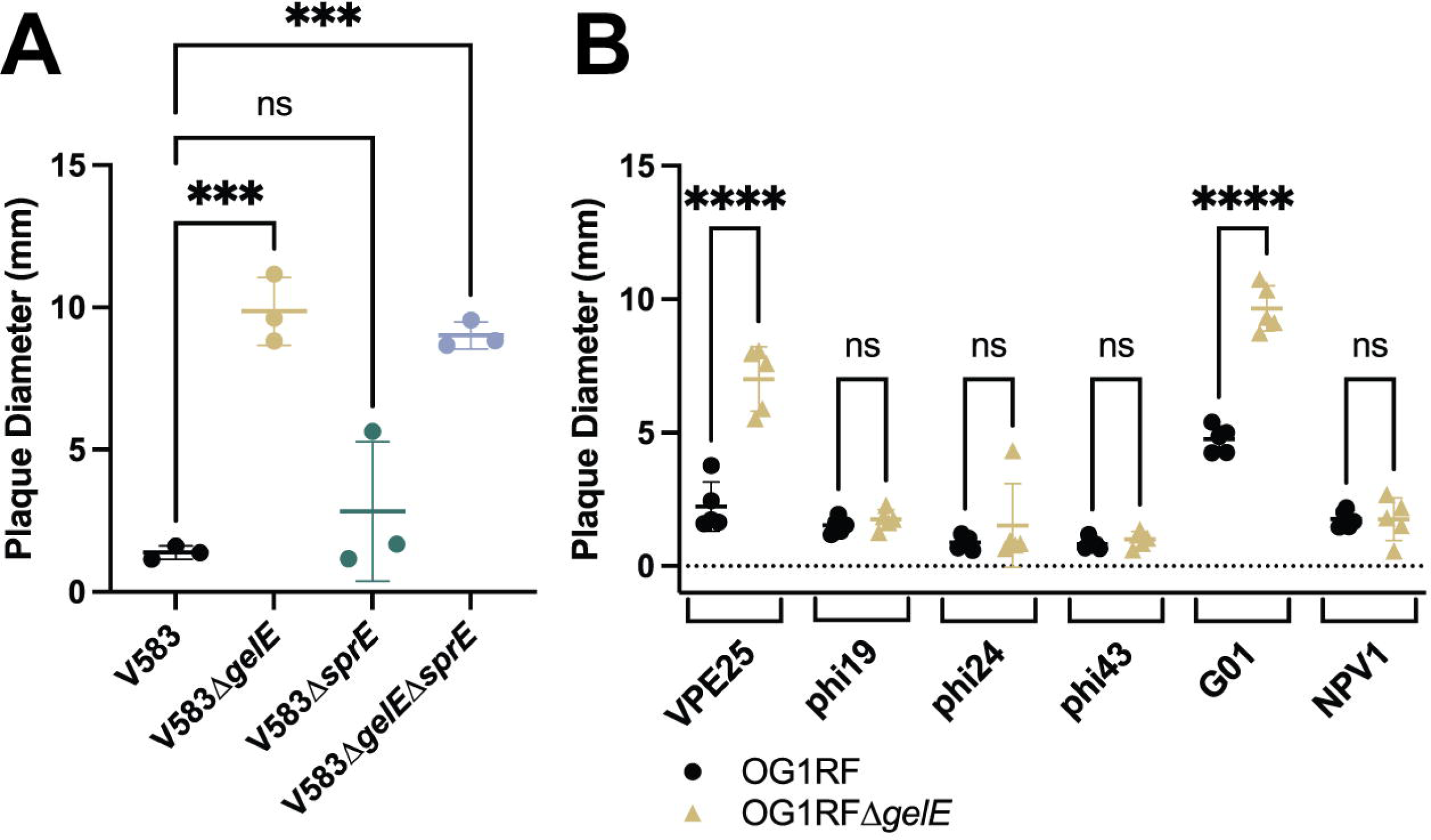
Infection by genetically distinct phages and bacterial species is impacted by the presence of GelE. **(A)** At 48 hours, average plaque diameter on V583Δ*gelE* and V583Δ*gelE*Δ*sprE* is significantly larger than wild type V583. Data represents average plaque diameter from three biological replicates. Significance determined using one-way ANOVA with multiple comparisons. **(B)** At 48 hours, the plaque diameter of VPE25 and G01 plaques is significantly larger on an overlay of OG1RFΔ*gelE* when compared to OG1RF. Data represents average plaque diameter from four biological replicates. Significance determined using two-way ANOVA with multiple comparisons. **(A,B)** Error bars represent standard deviation. **** p ≤ 0.0001, *** 0.0001 < p ≤ 0.001, ns = not significant.

To determine if this halo plaque morphology was unique to the phage VPE25, we used a panel of genetically distinct phages and measured plaque diameters on OG1RF and OG1RFΔ*gelE*. In addition to VPE25, the phage G01 produced a halo phenotype and significantly larger plaques after 48 hours of infection (Figure 3B). G01 is a 41 kbp phage isolated from the Ganges River in India (56). While both VPE25 and G01 are siphophages with icosahedral heads, the two phages share greater than 30% nucleotide identity in only four genes as identified via clinker (Supplemental Figure 4A) (81). Specifically, these homologous genes encode a lysin, glutaredoxin, a DUF1140 domain-containing protein, and a hypothetical protein. Interestingly, while the VPE25 and G01 lysins share 65% identity across 66% coverage, most of the homology is focused in the amidase domain (Supplemental Figure 4B) (82). These data indicate that GelE-mediated plaque morphology is neither *E. faecalis* strain- or phage-dependent.

### GelE regulation of the murein hydrolase regulator *lrgA* and antiholin-like protein *lrgB* provides protection from phage-mediated inhibition

Previous work in *E. faecalis* V583 has demonstrated an upregulation of the *lrgAB* operon in the presence of *gelE* (35). While the function of the *lrgAB* locus is uncharacterized in *E. faecalis*, LrgA and LrgB are involved in repression of murein hydrolase activity and decreased autolysis in *Staphylococcus aureus* and their expression is dependent on the Agr quorum-sensing system (83, 84). We hypothesized that *lrgA* and/or *lrgB* play a role in limiting phage-mediated haloed plaque formation. The *lrgAB* locus is regulated by the LytSR two-component system, located directly upstream of its operon. LytSR is hypothesized to activate in the presence of extracellular GelE, after which it increases transcription of *lrgAB* (35). While the functions of *lrgA* and *lrgB* have not been studied extensively in enterococci, they have been shown to be upregulated during growth in blood (85).

Considering the role of LrgA and LrgB in limiting murein hydrolase activity, we hypothesized that they may be involved in protection against phage-mediated extracellular lysis. VPE25 plaque diameters on *lytS*-Tn and *lytR*-Tn mutants were significantly larger than wild type (Figure 4A). Next, the plaque assay was repeated with *lrgA*-Tn and *lrgB*-Tn mutants. Data again show that plaque diameters are significantly larger than wild type OG1RF (Figure 4B). Thus, our data suggests that the murein hydrolase regulator LrgA and the anti-holin LrgB are ultimately responsible for the altered plaque morphology in the absence of GelE and supports the notion that this activity is mediated first by GelE and then by the LytSR two-component system. The underabundance of proteins like GelE during phage infection may be due in part to phage-mediated changes in enterococcal quorum-sensing to facilitate an environment more permissible to phage infection. This is further supported by the increased phage-dependent inhibition seen on plates in the absence of these genes.

**Figure 4:**
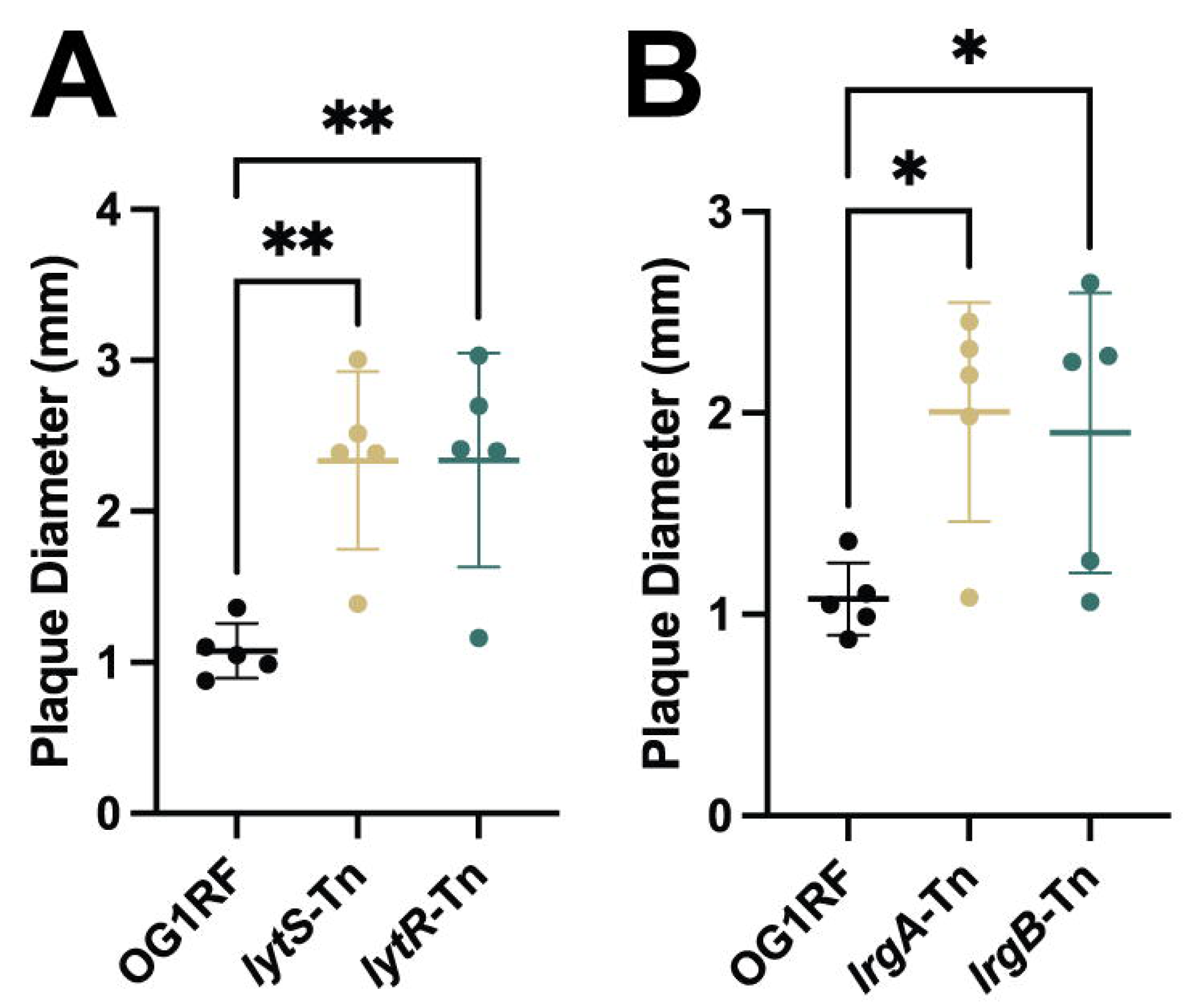
Mutation of *lrgA* and *lrgB* genes results in altered plaque morphology. **(A)** Average plaque diameter on *lytS*-Tn and *lytR*-Tn mutants is significantly larger than wild type. **(B)** Average plaque diameter on *lrgA*-Tn and *lrgB*-Tn mutants is significantly larger than wild type. **(A, B)** Data represents five biological replicates. Significance determined using one-way ANOVA with multiple comparisons. Error bars represent standard deviation. ** 0.001 < p ≤ 0.01, * 0.01 < p ≤ 0.05.

## DISCUSSION

The continued rise in antibiotic resistant bacterial infections has led to renewed interest in alternative antimicrobials, including phages. However, gaps in our understanding of phage biology and infection dynamics limit the full potential of emerging phage therapeutics. In this study, we used proteomics to probe the complexities of phage infection and understand the interplay between phages and *E. faecalis*. The resulting dataset provided insights in the timing of phage protein expression during infection, as well as identifying the altered abundance of hundreds of bacterial proteins during infection, primarily those involved in cell wall remodeling and bacterial metabolism. Additionally, this work revealed a role for the FsrA-regulated protease GelE, and to a lesser extent the protease SprE, in driving phenotypic changes in phage plaque morphology. This phenotypic change is due in part to ablated activity of the murein hydrolase regulator LrgA and the anti-holin LrgB, which we hypothesize aid in protection from increased phage-mediated attack (Figure 5). Future development of phages for clinical application may exploit this phenotype to allow for phage-mediated modulation of the virulence factor GelE to reduce *E. faecalis* virulence.

**Figure 5:**
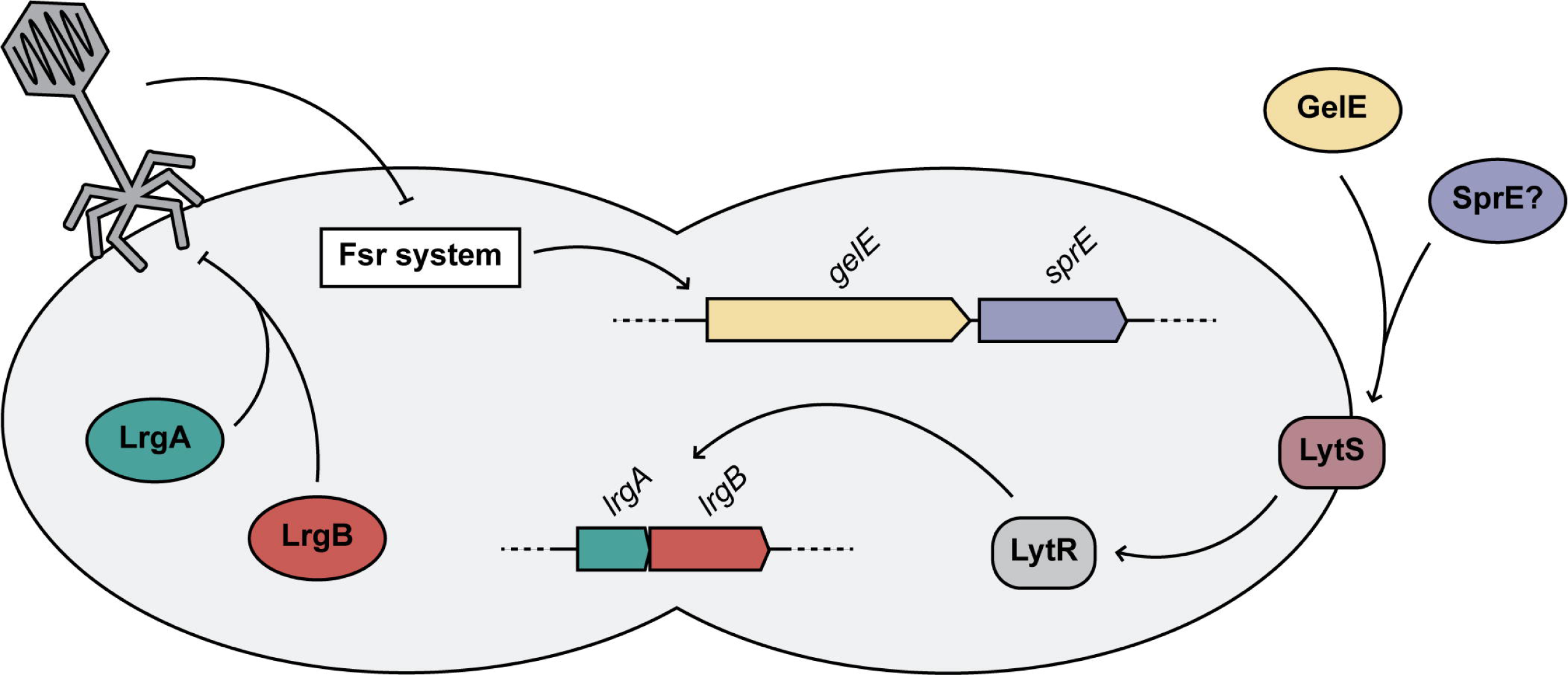
Enterococcal quorum-controlled proteases alter phage infection. Model figure summarizing the interactions between the Fsr quorum-sensing system, GelE, SprE, LrgA, LrgB and phage during infection of *E. faecalis*.

GelE was first investigated due to its significant under-abundance during phage infection (Figures 1C, 1D, Table 1, Supplemental Table 4). This observation is consistent with the repression of quorum-sensing-regulated transcripts during infection, including *gelE* and *sprE* (30). Gelatinase has been previously characterized as a virulence factor and has a role in the formation of biofilms, which are well-penetrated by phage (41, 42, 86–88). These data suggest that, in addition to aiding in the dispersal of biofilms, phage can limit biofilm initiation through repression of *gelE*. Further research may elucidate the phage factor responsible for mediating the repression of enterococcal quorum-sensing during infection. Inclusion of an identified gene or genes in a therapeutic phage may aid treatment by repressing the expression of *fsr*-regulated virulence factors (32, 45, 79).

While the absence of *gelE* is associated with the development of a haloed plaque in both *E. faecalis* OG1RF and V583, deletion of *sprE* results in changes to plaque morphology only in OG1RF (Figures 2C, 2E, 3A). These strains share 98.82% and 99.30% amino acid identity across the full sequences of GelE and SprE, respectively (82). With such high similarity and a well-characterized role for GelE in post-processing of SprE, it remains unclear why interruption of *sprE* does not result in haloed plaques in V583 (45). It is possible that SprE, in addition to GelE, is required for processing a downstream factor which is processed by GelE alone in V583, or in conjunction with an alternative protease. Post-processing by GelE is known to impact the localization of the autolysin AtlA on the cell surface (46). Bacteria often acquire phage resistance through mutations that result in changes to the structure of cell wall-associated molecules (54, 55). While the haloed plaques reported here are not dependent on AtlA, characterization of additional changes to the cell surface in the absence of *gelE*, *sprE*, or both can provide insight into the mechanism of the phenotypic change (Figure 2E).

Interruption of *lrgA* and *lrgB*, which are positively regulated in the presence of *gelE*, also resulted in the haloed plaque morphology (Figure 4B) (35). LrgA and LrgB are known to repress murein hydrolase activity and autolysis in *S. aureus*, suggesting a potential role for these proteins in limiting phage-mediated extracellular lysis in this model (83). However, the exact role of these proteins has not been studied in *Enterococcus*. Elucidation of the mechanism of LrgAB-conferred protection can inform our understanding of phage infection and enterococcal function. The similarity between the amidase domains of the G01 and VPE25 lysins, as well as the role of LrgA and LrgB in repression of murein hydrolase activity, suggests a possible protection against extracellular lysis conferred by the *lrgAB* operon (Figure 4B, Supplemental Figure 4B).

Similar haloed plaque morphologies have been associated with the presence of phage depolymerases (89, 90). These phage-encoded proteins can degrade a variety of bacterial polysaccharides, including capsular, extracellular, and lipopolysaccharides (91–93). While phage depolymerases have been previously associated with similar plaque phenotypes, analysis using InterProScan does not reveal any Pfam matches for such depolymerases within the genomes of phages VPE25 or G01 (73). While the mechanism of GelE-conferred protection from phage-mediated inhibition is still unknown, phages which are able to modulate the production of virulence factors such as GelE should be considered for their potential role in future therapeutic applications. Taken together, our work here provides an understanding of phage infection at the protein level and demonstrates a protective role for GelE, LrgA, and LrgB during phage infection (Figure 5).

## DATA AVAILABILITY

The proteomic data (MS raw files and search results) and the protein sequence database have been deposited to the ProteomeXchange Consortium via the PRIDE partner repository with the dataset identifier PXD026873 (94).

*Access for reviewers:* https://www.ebi.ac.uk/pride/archive/login*, Username, Password: TYHmjw0y)*.

## ACKNOWLEDGEMENTS

This work was supported by the National Institutes of Health grants R01AI141479 (B.A.D.) and R35GM138362 (M.K.). All LC-MS/MS measurements were made in the Molecular Education, Technology, and Research Innovation Centre (METRIC) at NC State University. The authors would like to thank Drs. Julia Willett, Kimberly Kline, and Danielle Garsin for providing bacterial strains used in this work.

## SUPPLEMENTAL FIGURE LEGENDS

**Supplemental Figure 1: Additional proteomic abundance changes and transposon screen. (A)** Volcano plot of bacterial proteins differentially abundant at 20 minutes post-infection between the infected and uninfected samples. Log2-transformed fold change between the two samples is measured on the x-axis. Negative log10-transformed q-value is measured on the y-axis with a line at a significance cut-off of q < 0.05. Points colored in blue are significantly underrepresented in the infected sample when compared to the uninfected sample. Points in red are significantly overrepresented. **(B)** Transposon mutant screen of phage infection indicates no changes in infectivity relative to OG1RF wild type. Transposon mutants were selected from the most over- or underrepresented proteins at 40 minutes as indicated by the differential abundance ratio (Table 1). Data represents an average of four replicates. Error bars represent standard deviation. Significance determined via one-way ANOVA with correction for multiple comparisons. ns = not significant.

**Supplemental Figure 2: Presence of GelE does not affect virion stability or adsorption. (A)** Azocoll cleavage indicates extracellular GelE activity peaks at 6 hours post-subculture in both the wild type strain OG1RF and the complement OG1RFΔ*gelE*::*gelE*. **(B)** Treatment of VPE25 virions with spent media has no effect on viable PFUs recovered, regardless of strain. **(B)** Adsorption of VPE25 to cells is not affected by strain. **(B,C)** Significance determined using one-way ANOVA with multiple comparisons. ns = not significant. **(A, B, C)** Data represents an average of three biological replicates. Error bars represent standard deviation.

**Supplemental Figure 3: Changes in plaque morphology on V583 are time-dependent.** At 24 hours, average VPE25 plaque diameter on V583Δ*gelE*, V583Δ*sprE*, and V583Δ*gelE*Δ*sprE* is not significantly different from wild type. Data represents three biological replicates. Significance determined using one-way ANOVA with multiple comparisons. Error bars represent standard deviation. ns = not significant.

**Supplemental Figure 4: VPE25 and G01 share significant homology in four genes, including their lysins. (A)** Comparison of phage genome similarities at the sequence level shows only four genes are conserved with >30% similarity between VPE25 and G01. Figure generated using clinker (81). **(B)** Schematic of VPE25 and G01 lysin homology generated via Blast 2 sequences (82). Matching amino acids are designated with the corresponding single-letter symbol in the middle lane. Conservative substitutions are marked with a “+”. Red text indicates residues in amidase domain, as identified via InterProScan (73).

## SUPPLEMENTAL TABLE LEGENDS

**Supplemental Table 1:** Bacterial strains, bacteriophages, plasmids, and primers used in this study

**Supplemental Table 2:** Phage average phage transcripts per million (TPM) at each timepoint from RNA-Seq transcriptomic reanalysis.

**Supplemental Table 3:** Phage proteins from mass spectrometry proteomic analysis

**Supplemental Table 4:** Bacterial proteins from mass spectrometry proteomic analysis

## Notes

### Competing Interest Statement

The authors have declared no competing interest.

