## Supplementary figures and images for "Enterococcal quorum-controlled protease alters phage infection"

### Supplementary Figure 1

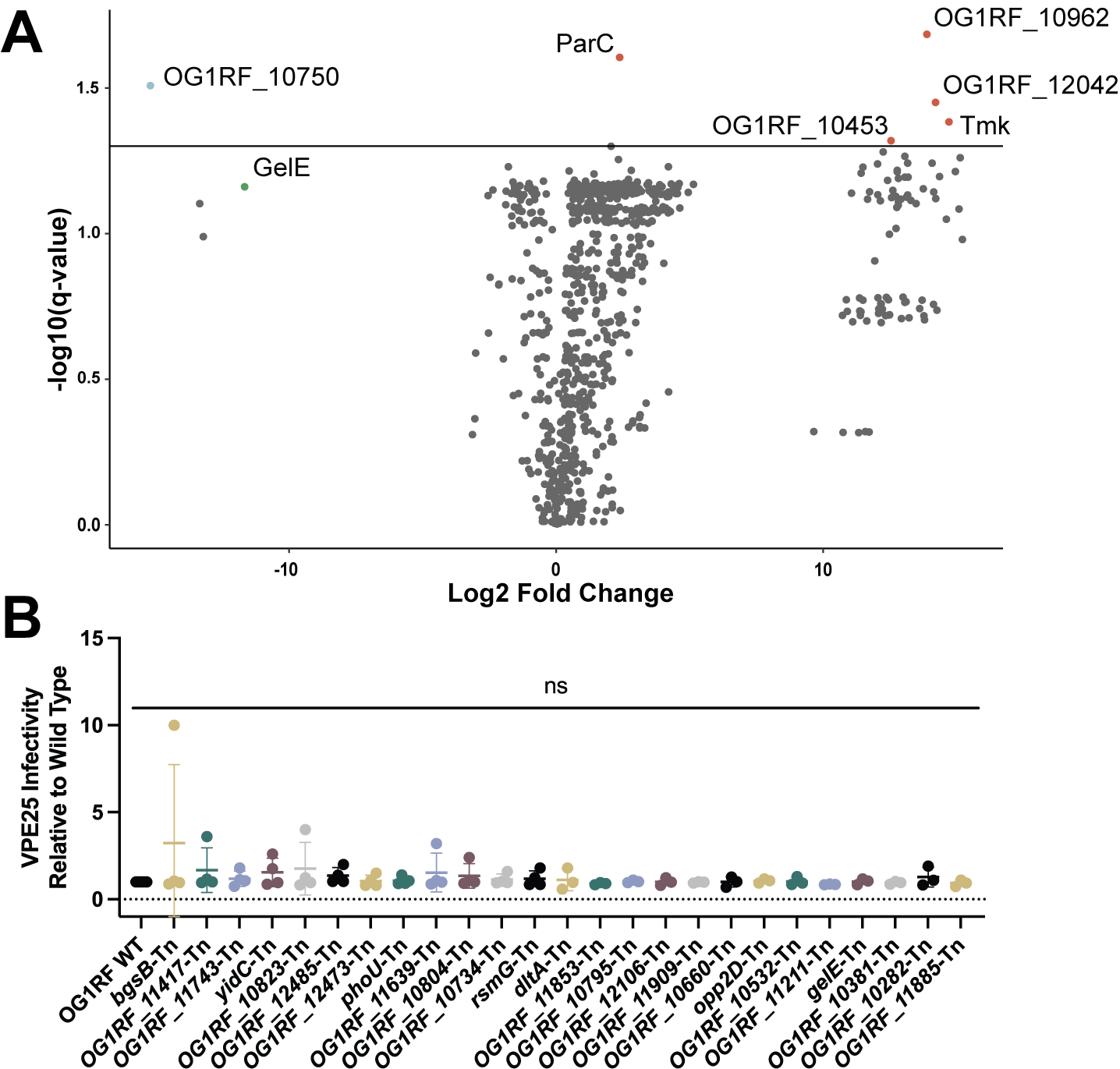

### Supplementary Figure 2

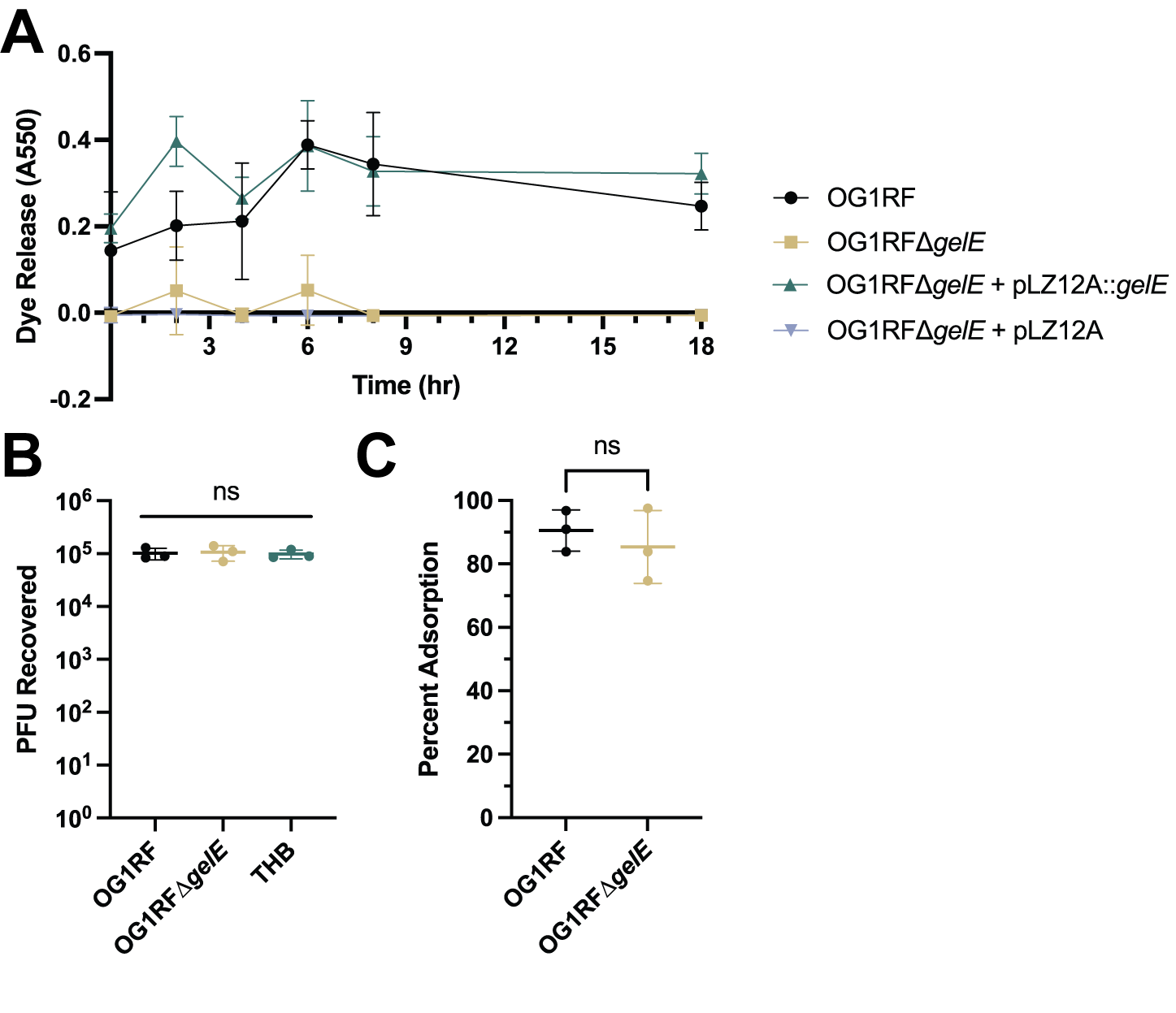

### Supplementary Figure 3

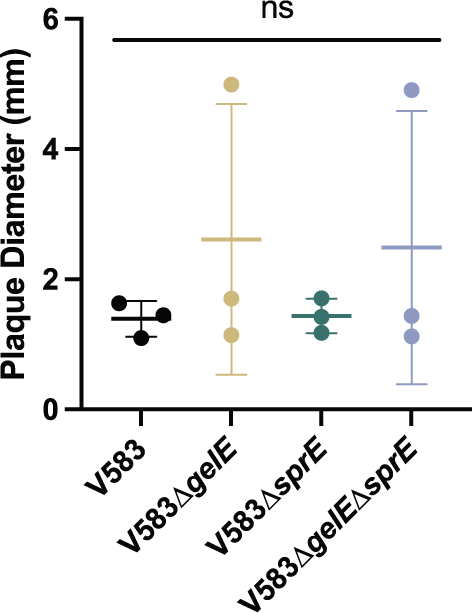

### Supplementary Figure 4

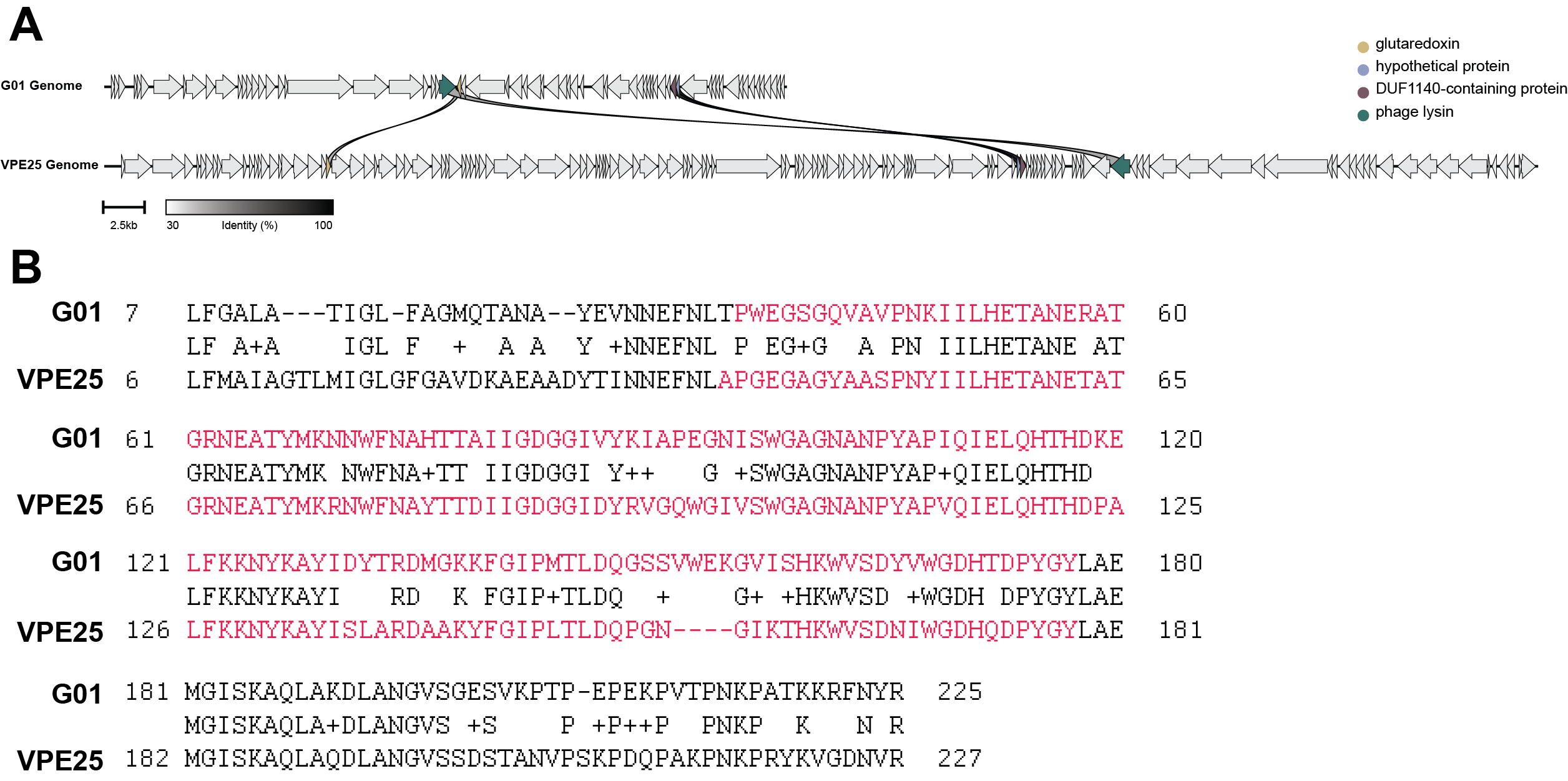
