## Supplementary Table 1 for "Enterococcal quorum-controlled protease alters phage infection"

**Supplemental Table 1**

| **Bacterial strain, phage, plasmid, or primer** | **Description** | **Reference** |
| --- | --- | --- |
| ***Enterococcus faecalis*** | | |
| OG1RF | Human oral isolate; Rf^R^, Fa^R^ | (1) |
| *bgsB*-Tn | *E. faecalis* OG1RF *bgsB* transposon mutant; Rf^R^, Fa^R^, Cm^R^ | (2) |
| *OG1RF_11417*-Tn | OG1RF *OG1RF_11417* transposon mutant; Rf^R^, Fa^R^, Cm^R^ | (2) |
| *OG1RF_11743*-Tn | OG1RF *OG1RF_11743* transposon mutant; Rf^R^, Fa^R^, Cm^R^ | (2) |
| *yidC*-Tn | OG1RF *yidC* transposon mutant; Rf^R^, Fa^R^, Cm^R^ | (2) |
| *OG1RF_10823*-Tn | OG1RF *OG1RF_10823* transposon mutant; Rf^R^, Fa^R^, Cm^R^ | (2) |
| *OG1RF_12485*-Tn | OG1RF *OG1RF_12485* transposon mutant; Rf^R^, Fa^R^, Cm^R^ | (2) |
| *OG1RF_12473*-Tn | OG1RF *OG1RF_12473* transposon mutant; Rf^R^, Fa^R^, Cm^R^ | (2) |
| *phoU*-Tn | OG1RF *phoU* transposon mutant; Rf^R^, Fa^R^, Cm^R^ | (2) |
| *OG1RF_11639*-Tn | OG1RF *OG1R_11639* transposon mutant; Rf^R^, Fa^R^, Cm^R^ | (2) |
| *OG1RF_10804*-Tn | OG1RF *OG1RF_10804* transposon mutant; Rf^R^, Fa^R^, Cm^R^ | (2) |
| *OG1RF_10734*-Tn | OG1RF *OG1RF_10734* transposon mutant; Rf^R^, Fa^R^, Cm^R^ | (2) |
| *rsmG*-Tn | OG1RF *rsmG* transposon mutant; Rf^R^, Fa^R^, Cm^R^ | (2) |
| *dltA*-Tn | OG1RF *dltA* transposon mutant; Rf^R^, Fa^R^, Cm^R^ | (2) |
| *OG1RF_11853*-Tn | OG1RF *OG1RF_11853* transposon mutant; Rf^R^, Fa^R^, Cm^R^ | (2) |
| *OG1RF_10795*-Tn | OG1RF *OG1RF_10795* transposon mutant; Rf^R^, Fa^R^, Cm^R^ | (2) |
| *OG1RF_12106*-Tn | OG1RF *OG1RF_12106* transposon mutant; Rf^R^, Fa^R^, Cm^R^ | (2) |
| *OG1RF_11909*-Tn | OG1RF *OG1RF_11909* transposon mutant; Rf^R^, Fa^R^, Cm^R^ | (2) |
| *OG1RF_10660*-Tn | OG1RF *OG1RF_10660* transposon mutant; Rf^R^, Fa^R^, Cm^R^ | (2) |
| *opp2D*-Tn | OG1RF *opp2D* transposon mutant; Rf^R^, Fa^R^, Cm^R^ | (2) |
| *OG1RF_10532*-Tn | OG1RF *OG1RF_10532* transposon mutant; Rf^R^, Fa^R^, Cm^R^ | (2) |
| *OG1RF_11211*-Tn | OG1RF *OG1RF_11211* transposon mutant; Rf^R^, Fa^R^, Cm^R^ | (2) |
| *gelE*-Tn | OG1RF *gelE* transposon mutant; Rf^R^, Fa^R^, Cm^R^ | (2) |
| *OG1RF_10381*-Tn | OG1RF *OG1RF_10381* transposon mutant; Rf^R^, Fa^R^, Cm^R^ | (2) |
| *OG1RF_10282*-Tn | OG1RF *OG1RF_10282* transposon mutant; Rf^R^, Fa^R^, Cm^R^ | (2) |
| *OG1RF_11885*-Tn | OG1RF *OG1RF_11885* transposon mutant; Rf^R^, Fa^R^, Cm^R^ | (2) |
| OG1RF∆*gelE* | OG1RF internal in-frame deletion in *gelE*, also called TX5264; Rf^R^, Fa^R^ | (3) |
| OG1RF∆*gelE*::*gelE* | OG1RF∆*gelE* complemented with *gelE* via pLZ12A::*gelE*; Rf^R^, Fa^R^, Cm^R^ | This study |
| OG1RF∆*gelE*::EV | OG1RF∆*gelE* carrying pLZ12A empty vector; Rf^R^, Fa^R^, Cm^R^ | This study |
| OG1RF∆*fsrA* | OG1RF with isogenic deletion of *fsrA*, also called JD100; Rf^R^, Fa^R^ | (4) |
| OG1RF∆*entV* | OG1RF with markerless deletion of *entV*, also called CEGF1; Rf^R^, Fa^R^ | (5) |
| OG1RF_*sprE*-Tn | OG1RF *sprE* transposon mutant; Rf^R^, Fa^R^, Cm^R^ | (2) |
| OG1RF∆*atlA* | OG1RF markerless deletion of *atlA*, also called OG1RF∆*0799*; Rf^R^, Fa^R^ | (6) |
| V583 | Human blood isolate; Vm^R^, Em^R^, Gm^R^ | (7) |
| V583∆*gelE* | V583 with isogenic markerless deletion of *gelE*, also called VT01; Vm^R^, Em^R^, Gm^R^ | (8) |
| V583∆*sprE* | V583 with isogenic markerless deletion of *sprE*, also called VT02; Vm^R^, Em^R^, Gm^R^ | (8) |
| V583∆*gelE*∆*sprE* | V583 with isogenic markerless deletion of *gelE* and *sprE*, also called VT03; Vm^R^, Em^R^, Gm^R^ | (8) |
| *lrgA*-Tn | OG1RF *lrgA* transposon mutant with insertion at position 2,597,437; Rf^R^, Fa^R^, Cm^R^ | (2) |
| *lrgB*-Tn | OG1RF *lrgB* transposon mutant with insertion at position 2,597,095; Rf^R^, Fa^R^, Cm^R^ | (2) |
| *lytS*-Tn | OG1RF *lytS* transposon mutant with insertion at position 2,600,097; Rf^R^, Fa^R^, Cm^R^ | (2) |
| *lytR*-Tn | OG1RF *lytR* transposon mutant with insertion at position 2,598,216; Rf^R^, Fa^R^, Cm^R^ | (2) |
| **Bacteriophages** | | |
| VPE25 | Wastewater isolate | (9) |
| phi19 | Obtained from Naval Medical Research Center (NMRC) | (10) |
| phi24 | Obtained from NMRC | (10) |
| phi43 | Obtained from NMRC | (10) |
| G01 | Ganges River water isolate | (11) |
| NPV1 | Wastewater isolate | (12) |
| **Plasmids** | | |
| pLZ12A | *bacA* promoter cloned into shuttle vector pLZ12; pSH71 origin; Cm^R^ | (10, 13) |
| pLZ12A::*gelE* | pLZ12A expressing *gelE* from P_bacA_; Cloned into PstI/BamHI site; Cm^R^ | This study |
| pLZ12A::*lrgAB* | pLZ12A expressing *lrgA* and *lrgB* from P_bacA_; Cloned into EcoRI/BamHI site; Cm^R^ | This study |
| **Primers** | | |
| 526insert_fwd | NNNNNNCTGCAGTTGATGAAGGGAAATAAAATTTTATACATT; forward primer to generate pLZ12A::*gelE*, PstI site | This study |
| 526insert_rev | NNNNNNGGATCCTCATTCATTGACCAGAACAGATT; reverse primer to generate pLZ12A::*gelE*; BamHI site | This study |
| lrgAinsert_fwd | NNNNNNGAATTCATGGAAAAGAAAGTATATTCATTTTTACAA; forward primer to generate pLZ12A::*lrgAB*, EcoRI site | This study |
| lrgBinsert_rev | NNNNNNGGATCCTTATAGACCAATGACCGTTGCA; reverse primer to generate pLZ12A::*lrgAB*, BamHI site | This study |
| 12192tn_fwd | CATGACACCCGTTCCAATAAAG; forward primer to confirm *bgsB*-Tn transposon insertion | This study |
| 12192tn_rev | AGAAGGGAACGCCTATTTAAATAATT; reverse primer to confirm *bgsB*-Tn transposon insertion | This study |
| 11417tn_fwd | GTGTCGCCATGTAATGTGC; forward primer to confirm *OG1RF_11417*-Tn transposon insertion | This study |
| 11417tn_rev | ACAGCGGAGATCATTAAAGTTCA; reverse primer to confirm *OG1RF_11417*-Tn transposon insertion | This study |
| 11743tn_fwd | AGGTTACACTTTTTTTAATTTTTATAGGTC; forward primer to confirm *OG1RF_11743*-Tn transposon insertion | This study |
| 11743tn_rev | CTTAGCGAAAGAAGTCTTTCCAG; reverse primer to confirm *OG1RF_11743*-Tn transposon insertion | This study |
| 11837tn_fwd | AATTGGAACTATGGTTGGGCG; forward primer to confirm *yidC*-Tn transposon insertion | This study |
| 11837tn_rev | CTAAATAGGCTAAACCAGCTAATGC; reverse primer to confirm *yidC*-Tn transposon insertion | This study |
| 10823tn_fwd | TGAAAGTTTTATCGTAGAAGGTCC; forward primer to confirm *OG1RF_10823*-Tn transposon insertion | This study |
| 10823tn_rev | TTCGTACCACTCTTTCTGAATTG; reverse primer to confirm *OG1RF_10823*-Tn transposon insertion | This study |
| 12485tn_fwd | GTAAGATCTTTTTCTTCTGTCGTTTTATT; forward primer to confirm *OG1RF_12485*-Tn transposon insertion | This study |
| 12485tn_rev | ATCAAGCAACGCAGAAATTTATTG; reverse primer to confirm *OG1RF_12485*-Tn transposon insertion | This study |
| 12473tn_fwd | ACACTAACAGGTAAGCCACC; forward primer to confirm *OG1RF_12473*-Tn transposon insertion | This study |
| 12473tn_rev | TGCCAAAGCCCTTTTTTGTT; reverse primer to confirm *OG1RF_12473*-Tn transposon insertion | This study |
| 11465tn_fwd | AAAATCAGTTCATTTTACGCTAAAAAAAC; forward primer to confirm *phoU*-Tn transposon insertion | This study |
| 11465tn_rev | AAATTTCAGATATGTCTGATTATGTGAAAA; reverse primer to confirm *phoU*-Tn transposon insertion | This study |
| 11639tn_fwd | CAATCCATTGTTCTTTTTCTGTTTCTG; forward primer to confirm *OG1RF_11639*-Tn transposon insertion | This study |
| 11639tn_rev | GGCCTTGATTGGGATCGTAAA; reverse primer to confirm *OG1RF_11639*-Tn transposon insertion | This study |
| 10804tn_fwd | TATCATGAAAATAAAGCCAAATTATTACAT; forward primer to confirm *OG1RF_10804*-Tn transposon insertion | This study |
| 10804tn_rev | GCTCTTCTGTATGTTGCGGA; reverse primer to confirm *OG1RF_10804*-Tn transposon insertion | This study |
| 10734tn_fwd | ACATCCTTTTATTGACACTGTCGG; forward primer to confirm *OG1RF_10734*-Tn transposon insertion | This study |
| 10734tn_rev | GTTAACAACGAAGCCAGCTACT; reverse primer to confirm *OG1RF_10734*-Tn transposon insertion | This study |
| 12545tn_fwd | ACATCTGGTTTAGGAACACCC; forward primer to confirm *rsmG*-Tn transposon insertion | This study |
| 12545tn_rev | GCTATTTTTTAGCCTTAAAAGCATCG; reverse primer to confirm *rsmG*-Tn transposon insertion | This study |
| 12112tn_fwd | GTGTTTGGTACCAAGGCCAA; forward primer to confirm *dltA*-Tn transposon insertion | This study |
| 12112tn_rev | GGATCATCACTTAACGAATGTTTCC; reverse primer to confirm *dltA*-Tn transposon insertion | This study |
| new_11853tn_fwd | CAATCCGCGAATTCCACGAATA; forward primer to confirm *OG1RF_11853*-Tn transposon insertion | This study |
| new_11853_rev | ACATATTGTGACTGCCAATAAAGATTTAC; reverse primer to confirm *OG1RF_11853*-Tn transposon insertion | This study |
| 10795tn_fwd | TCAATTTATTTCAATCAATGTATGGATCTC; forward primer to confirm *OG1RF_10795*-Tn transposon insertion | This study |
| 10795tn_rev | AATTCTAACTGTTGCCTGATATCATC; reverse primer to confirm *OG1RF_10795*-Tn transposon insertion | This study |
| 12106tn_fwd | GCGCTTCTTCCCGTAGTT; forward primer to confirm *OG1RF_12106*-Tn transposon insertion | This study |
| 12106tn_rev | ACACAAGATCAAATGCATTTCAAAG; reverse primer to confirm *OG1RF_12106*-Tn transposon insertion | This study |
| 11909tn_fwd | TTGTCAAAAAACGTCCAACAAAAAA; forward primer to confirm *OG1RF_11909*-Tn transposon insertion | This study |
| 11909tn_rev | CTAACGTTGCTAGTCCTAAACGG; reverse primer to confirm *OG1RF_11909*-Tn transposon insertion | This study |
| 10660tn_fwd | TTAAAAAGCAGTGCCTACTCGTC; forward primer to confirm *OG1RF_10660*-Tn transposon insertion | This study |
| 10660tn_rev | AATCGTCACTTCCATTGCAACT; reverse primer to confirm *OG1RF_10660*-Tn transposon insertion | This study |
| 12370tn_fwd | GGTAATTTGGTTAACGAAGGAACAA; forward primer to confirm *opp2D*-Tn transposon insertion | This study |
| 12370tn_rev | AACAGTATCCTCATGAATTGTCTGG; reverse primer to confirm *opp2D*-Tn transposon insertion | This study |
| 10532tn_fwd | GGCGCCAGATCCACCAATTA; forward primer to confirm *OG1RF_10532*-Tn transposon insertion | This study |
| 10532tn_rev | CACAGCATCGGCCGCTAA; reverse primer to confirm *OG1RF_10532*-Tn transposon insertion | This study |
| 11211tn_fwd | CATTCATACAAAAAACTCTTCCG; forward primer to confirm *OG1RF_11211*-Tn transposon insertion | This study |
| 11211tn_rev | GGTTTCTTTATTGATAATCTTTTCTAATAG; reverse primer to confirm *OG1RF_11211*-Tn transposon insertion | This study |
| 11526tn_fwd | TGTTTACTTGGCGTTTGTAAATTTC; forward primer to confirm *gelE*-Tn transposon insertion | This study |
| 11526tn_rev | CGCATGGAAAATTTGTGAAAACA; reverse primer to confirm *gelE*-Tn transposon insertion | This study |
| 10381tn_fwd | TGGAACAAACATCCAAAGTGATTG; forward primer to confirm *OG1RF_10381*-Tn transposon insertion | This study |
| 10381tn_rev | AAACTAAAGCTGACATTCCCGA; reverse primer to confirm *OG1RF_10381*-Tn transposon insertion | This study |
| 10282tn_fwd | GTAACCACACACATTTGCCG; forward primer to confirm *OG1RF_10282*-Tn transposon insertion | This study |
| 10282tn_rev | TTTTTTACCATACTTCTCTAGCAATTTC; reverse primer to confirm *OG1RF_10282*-Tn transposon insertion | This study |
| 11885tn_fwd | GTCGCCATCTAACGCTTTCA; forward primer to confirm *OG1RF_11885*-Tn transposon insertion | This study |
| 11885tn_rev | TTCACTATGTTTCACGAAAAATGAAATATG; reverse primer to confirm *OG1RF_11885*-Tn transposon insertion | This study |
| 11525tn_fwd | CTGCTGGCACAGCGGATAAA; forward primer to confirm *sprE*-Tn transposon insertion | This study |
| 11525tn_rev | GTGACTGTCGGCAAACAAAACG; reverse primer to confirm *sprE*-Tn transposon insertion | This study |
| lrgAtn_fwd | CCCTAAAGAATTAACTAAGGAAATCCC; forward primer to confirm *lrgA*-Tn transposon insertion | This study |
| lrgAtn_rev | GCCATTATTATGTTAATCGCCAAC; reverse primer to confirm *lrgA*-Tn transposon insertion | This study |
| lrgBtn_fwd | GCCACCTTCATTGTATTGCTC; forward primer to confirm *lrgB*-Tn transposon insertion | This study |
| lrgBtn_rev | CCCGCAGCAACGAAGAA; reverse primer to confirm *lrgB*-Tn transposon insertion | This study |
| lytRtn_fwd | GGTCCTTTTTCGGCAGTGT; forward primer to confirm *lytR*-Tn transposon insertion | This study |
| lytRtn_rev | GCACGTATTAATCGTTGATGATGAG; forward primer to confirm *lytR*-Tn transposon insertion | This study |
| lytStn_fwd | GGGAACAATCTGATTCTTAGCGATT; forward primer to confirm *lytS*-Tn transposon insertion | This study |
| lytStn_rev | CTAAAGGCGCTAATATTCCTGCG; forward primer to confirm *lytS*-Tn transposon insertion | This study |

1.	Bourgogne A, Garsin DA, Qin X, Singh KV, Sillanpaa J, Yerrapragada S, Ding Y, Dugan-Rocha S, Buhay C, Shen H, Chen G, Williams G, Muzny D, Maadani A, Fox KA, Gioia J, Chen L, Shang Y, Arias CA, Nallapareddy SR, Zhao M, Prakash VP, Chowdhury S, Jiang H, Gibbs RA, Murray BE, Highlander SK, Weinstock GM. 2008. Large scale variation in Enterococcus faecalis illustrated by the genome analysis of strain OG1RF. Genome Biology 9:R110.

2.	Dale JL, Beckman KB, Willett JLE, Nilson JL, Palani NP, Baller JA, Hauge A, Gohl DM, Erickson R, Manias DA, Sadowsky MJ, Dunny GM. 2018. Comprehensive functional analysis of the Enterococcus faecalis core genome using an ordered, sequence-defined collection of insertional mutations in strain OG1RF. mSystems 3.

3.	Qin X. 2000. Identification and characterization of virulence in *Enterococcus faecalis*. Ph.D. University of Texas Graduate School of Biomedical Sciences, Houston.

9.	Duerkop BA, Huo W, Bhardwaj P, Palmer KL, Hooper LV. 2016. Molecular basis for lytic bacteriophage resistance in enterococci. mBio 7.

10.	Chatterjee A, Johnson CN, Luong P, Hullahalli K, McBride SW, Schubert AM, Palmer KL, Carlson PE, Jr., Duerkop BA. 2019. Bacteriophage resistance alters antibiotic-mediated intestinal expansion of enterococci. Infect Immun 87.

11.	Sheriff EK, Andersen SE, Chatterjee A, Duerkop BA. 2024. Complete genome sequence of enterococcal phage G01. Microbiol Resour Announc doi:10.1128/mra.01217-23:e0121723.

12.	Trotter KM, Dunny GM. 1990. Mutants of Enterococcus faecalis deficient as recipients in mating with donors carrying pheromone-inducible plasmids. Plasmid 24:57-67.

13.	Perez-Casal J, Caparon MG, Scott JR. 1991. Mry, a trans-acting positive regulator of the M protein gene of Streptococcus pyogenes with similarity to the receptor proteins of two-component regulatory systems. J Bacteriol 173:2617-24.
