## Supplementary Methods for "Enterococcal quorum-controlled protease alters phage infection"

**Azocoll assays**

Azocoll assays were performed as previously described with slight modification (46, 47). 0.125 g Azocoll (Azo dye-impregnated collagen, Sigma-Aldrich) was added to 25 mL KPBS on a plate rotor for 1 hr. Azocoll was then pelleted via centrifugation at 4,600 × g for 10 minutes. Supernatant was removed, and wash was repeated. Washed Azocoll was resuspended in 25 mL KPBS supplemented with 1 mM CaCl_2_. Overnight bacterial cultures were diluted to an OD_600_ of 0.025 and grown at 37°C with aeration for the time indicated. After appropriate growth, cells were pelleted at 21,000 × g for 1 minute. 500 µL of supernatant was mixed with 500 µL washed Azocoll and incubated overnight at 37°C with aeration. The following day, remaining Azocoll substrate was pelleted via centrifugation at 21,000 × g for 1 minute. A550 of the supernatant was measured to determine absolute gelatinase activity.

**Spent media treatment of phage virions**

Overnight bacterial cultures were subcultured to an OD_600_ of 0.025. Subcultures were incubated for 6 hours at 37°C with aeration to encourage peak GelE activity, as measured via Azocoll assay (Supplemental Figure 2A). Following incubation, cultures were spun down at 4,600 × g for 3 minutes, and supernatant was 0.45 µm filter sterilized. 10^5^ PFU of VPE25 was added to 100 µL of spent media and mixed gently by inversion. Plain THB was used as a negative control. Treatments were incubated for 18 hours at room temperature before plating on OG1RF top agar overlays to enumerate PFU.

**Adsorption assays**

Adsorption assays were performed as previously described with slight modifications (43, 44). Overnight cultures of bacteria were pelleted and resuspended in SM+ buffer. Cultures were then normalized to an OD_600_ of 0.1. 500 µL of dilute overnight culture was added to VPE25 at an MOI of 0.1 and mixed gently by inversion. Phage were allowed to adsorb for 15 minutes before bacterial cells were spun out at 21,000 × g for 2 minutes. Remaining phages in supernatant were enumerated via plaque assay. SM+ buffer was used as a no-adsorption control. Percent adsorption was calculated as $\frac{PFU\left( control \right)-PFU(supernatant)}{PFU(control)}\times100\%$.
